# EBD-DTI: Episodic Bridge Diffusion for Zero-Shot Cold-Start Drug-Target Interaction Prediction

**DOI:** 10.64898/2026.07.14.738384

**Authors:** Jiongxin Liu, Jiameng Le, Chuanru Wei, Mingming Liu, Zixuan Yin

## Abstract

Predicting drug–target interactions (DTI) for entirely unseen drugs or proteins—the cold-start problem—remains a critical challenge in computational drug discovery. While sequence-based methods naturally support zero-shot generalization, they often ignore relational topology, and existing graph-based approaches either rely on global diffusion that blurs the boundary between inductive and transductive evaluation or require a few known interaction samples at test time (few-shot). We present EBD-DTI, a framework that enables zero-shot inference in graph-based DTI models without requiring any known interactions for unseen entities. The key innovation is *episodic cold-start training* : at each epoch, a random subset of training entities is masked and treated as pseudo-cold, forcing the model to learn cold-start inference with explicit gradient supervision. A bridge-conditioned local subgraph, together with multi-hop diffusion, provides cold entities with relational context from their nearest observed neighbors. Experiments on three benchmarks (BioSNAP, BindingDB, and DrugBank) demonstrate that EBD-DTI achieves competitive or superior performance compared to state-of-the-art methods under strict zero-shot evaluation, with episodic training improving AUC by up to 12%.

## 1 Introduction

Identifying which drugs bind which proteins is central to computational drug discovery: it lets researchers narrow down candidates before committing to expensive wet-lab screening across millions of compounds [22]. When a new chemical entity is synthesized or a previously unstudied protein is characterized, predicting its interactions becomes a critical bottleneck: experimental validation is time-consuming, expensive, and infeasible at scale. The ability to computationally infer interactions for unseen molecules and proteins could accelerate discovery timelines by months or years, and is especially valuable for emerging disease targets where no historical interaction data exists.

Recent methods have driven substantial progress in DTI prediction, ranging from graph neural networks such as GCN [9] and GAT [17] that propagate information through interaction graphs, to sequence-based models such as DrugBAN [1] that encode drug–protein pairs directly from molecular features. These methods achieve strong performance when drugs and proteins appear in the training data, leveraging known interaction structure or learned feature representations as inductive signals.

However, this progress obscures a fundamental limitation: these methods operate in the transductive cold-start regime, where the model has never learned to predict interactions for drugs or proteins that are entirely absent from training data. In real-world discovery, newly synthesized compounds and recently characterized proteins have zero prior interaction records and therefore zero connectivity in the training graph. Standard GNN message passing cannot handle such entities by definition—information propagates through edges, and absent entities have no edges.

The gap shows up clearly in practice. Under our strict zero-shot protocol, DrugBAN [1] achieves AUC=0.873 on cold-drug but drops to 0.764 on cold-target ( *−*12.5% on BioSNAP); HyperAttentionDTI [24] drops from AUC=0.957 (cold-drug) to 0.689 (cold-target) on BindingDB ( *−*28.0%). These sharp directional asymmetries suggest that protein representations generalize poorly to unseen targets: models can handle new drugs reasonably well (seen proteins provide useful context), but collapse when confronted with a new protein whose interaction pattern was never observed during training.

Graph propagation methods such as APPNP [10] can be adapted for cold-start by constructing KNN edges for unseen entities in post-hoc fashion, but they introduce a subtle but critical problem: the KNN edges are added to the evaluation graph and remain fixed, meaning test entities gain access to structure during inference that was predetermined at training time. This creates a setting that blurs the boundary between inductive and transductive evaluation, as test entities gain access to pre-defined structural context. In genuinely cold-start scenarios—where entities are completely novel and no prior information is available— these methods cannot be deployed without test-time graph modification.

An alternative is meta-learning: methods such as ZeroBind [18], MGDTI [19], and BioBridge [12] use few-shot adaptation or prototypical networks, allowing the model to learn from a small support set of known interactions for a cold entity at test time. While conceptually elegant, this approach assumes that test-time support samples are available—a requirement that often fails in practice. For a newly discovered protein without any known interactions, or in time-critical drug discovery pipelines where experimental validation cannot be performed, such samples do not exist.

We identify a more fundamental problem underlying all transductive approaches: the bridge mechanism that connects cold entities to the training graph never receives gradient signal during training, because all training entities are “seen.” In standard supervised DTI learning, the model learns to classify interactions for known entities; the bridge is never invoked, and therefore never learns to select appropriate neighbors for unseen entities. When deployed on test data with genuinely cold entities, the bridge operates at random initialization—its parameters have never been exposed to a signal that teaches it how to generalize. From a domain adaptation perspective [2], this reflects a profound training-test distribution mismatch: the training distribution has all entities observed; the test distribution has cold entities unobserved. The model has learned a function that assumes the world of training, not the world of testing.

To address this signal disconnection and distribution mismatch, we propose EBD-DTI (Episodic Bridge Diffusion for Cold-Start DTI), which introduces a novel training mechanism aligned with the test-time cold-start challenge. Rather than training on seen entities alone and hoping the bridge generalizes, EBD-DTI explicitly simulates cold-start scenarios during training through episodic masking: each epoch, 20% of training entities are randomly masked, forcing the bridge module to retrieve neighbors and propagate information from partial graphs, exactly as it must at test time. This episodic training provides supervised gradient signal directly to the bridge, teaching it to select appropriate neighbors and interpret their interaction patterns for prediction.

EBD-DTI combines this episodic training with two complementary technical innovations: (1) *bridge-conditioned local subgraph construction*, which dynamically creates an instance-specific local interaction neigh-borhood for each cold entity (distinct from global KNN graphs, eliminating semi-transductive information leakage); and (2) *instance-specific local diffusion*, which propagates information through the neighborhood in T=3 steps, requiring substantially fewer diffusion steps than global methods while respecting the DTI-structure of interaction networks. Unlike global diffusion methods that embed test entities into a pre-computed global structure, unlike fixed KNN graphs that determine test neighborhoods at training time, and unlike sequence-only baselines that lack explicit cold-start training signal, EBD-DTI uses episodic training to directly teach the model how to generalize to unseen entities.

Our contributions are:

1. **Episodic cold-start training mechanism**. We propose a training strategy that directly addresses the training-test distribution mismatch by randomly masking training entities each epoch to create pseudo-cold scenarios, providing supervised gradient signal to the bridge module. This is the first method to explicitly train a bridge mechanism under cold-start conditions, closing the generalization gap that causes other methods to fail.
2. **Dynamic bridge-conditioned local subgraph construction**. For each cold entity, we construct a distinct instance-specific subgraph centered on its feature-space neighbors, enabling strict zero-shot inference without assuming test-time support sets or pre-computed graph structure.
3. **Comprehensive strict zero-shot evaluation**. We evaluate on three benchmarks (BioSNAP, BindingDB, DrugBank) under truly zero-shot cold-start protocols where all test-entity edges are completely removed. EBD-DTI achieves leading cold-target AUPR on BioSNAP (0.919) and competitive or superior results across benchmarks, with episodic training improving cold-target AUC by up to 12%. Ablation studies confirm that episodic training drives the improvement, not the graph architecture.

## 2 Related Work

*Graph-Based DTI Methods*. Graph neural networks such as GCN, GAT, and GraphSAGE learn node representations by aggregating information from local neighborhoods through message passing. Recent extensions incorporate more expressive architectures: DrugBAN [1] introduces bilinear attention, while MolTrans and HyperAttentionDTI employ transformer-based encoders. Despite their strong empirical performance, these methods fundamentally operate in a transductive setting, where all test entities are assumed to be embedded within the training graph. As a result, they cannot naturally handle entities with no observed interactions. In contrast, EBD-DTI performs inference on *dynamically constructed local subgraphs* tailored to each cold entity, enabling prediction without requiring pre-existing graph connectivity.

### Cold-Start DTI

Cold-start DTI aims to generalize to entirely unseen drugs or proteins, a setting that more closely reflects real-world discovery scenarios. Recent methods such as ColdstartCPI [23] and SCOPEDTI [3] incorporate pre-trained representations or semi-inductive evaluation protocols to improve generalization. However, existing approaches typically rely on either auxiliary information (e.g., pre-trained embeddings) or evaluation protocols in which test entities are structurally incorporated into the graph prior to inference [15]. This implicitly assumes access to test-time structure, weakening the strict cold-start assumption. In contrast, EBD-DTI adopts a strictly zero-shot protocol in which cold entities have *no edges* in the evaluation graph, and directly addresses the missing training signal for cold-start inference through episodic simulation.

### Meta-Learning for DTI

Meta-learning approaches such as ZeroBind, MGDTI, and BioBridge formulate DTI prediction as a few-shot learning problem, enabling adaptation to new entities using a small support set of known interactions. While effective under this assumption, their applicability is limited in scenarios where no interaction data is available for new entities. EBD-DTI removes this requirement entirely by performing zero-shot inference, making it applicable to truly novel drugs and proteins without any test-time supervision.

*Masked Training*. Masked modeling has been widely used for representation learning, such as token masking in BERT [4] and node feature masking in GraphMAE [6]. However, these approaches focus on self-supervised reconstruction objectives and do not address cold-start generalization. Our episodic training repurposes masking in a fundamentally different way: instead of masking features, we mask entire *entities* to simulate cold-start conditions, while retaining supervised DTI labels. This provides explicit gradient signal to train the bridge mechanism, enabling the model to learn how to connect and reason over unseen entities.

## 3. Method

### 3.1 Problem Formulation

Given a set of drugs *D* = {*d*_1_, …, *d*_*M*_*}* and proteins *P* = {*p*_1_, …, *p*_*N*_*}*, together with known interactions *Y ⊆ D × P*, the goal is to predict whether an unknown pair (*d*_*i*_, *p*_*j*_) interacts. In the **cold-drug** setting, test drugs are entirely absent from training: *D* _test_ *∩ D*_train_ = ∅. In the **cold-target** setting, *P*_test_ ∩ *P*_train_ = ∅. We adopt a *strictly zero-shot* protocol: no known interactions involving cold entities are available at test time, unlike few-shot methods [12, 18].

### 3.2 Core Insights

Cold-start DTI prediction requires bridging the gap between training (all entities observed) and testing (entities unseen). We highlight three design insights underlying EBD-DTI.

#### Episodic training provides missing supervision

In standard supervised learning, all training entities are observed, so the bridge mechanism never receives gradient signal from cold-start scenarios. Episodic training addresses this by masking a random subset of entities each epoch, forcing the model to perform cold-start inference under real supervision.

#### Instance-specific subgraphs avoid structural bias

Cold entities lack graph connectivity and must obtain relational context dynamically. EBD-DTI constructs an instance-specific local subgraph for each cold entity based on feature-space neighbors, ensuring inference depends only on training data and the current model state—not on pre-defined graph structures.

#### Local diffusion matches DTI structure

DTI graphs are bipartite with small effective diameter; relevant context is typically reachable within a few hops. Local diffusion on instance-specific subgraphs captures this efficiently, avoiding the complexity of global diffusion.

### 3.3 Model Overview

Figure 1 illustrates the overall EBD-DTI pipeline.

**Figure 1.**
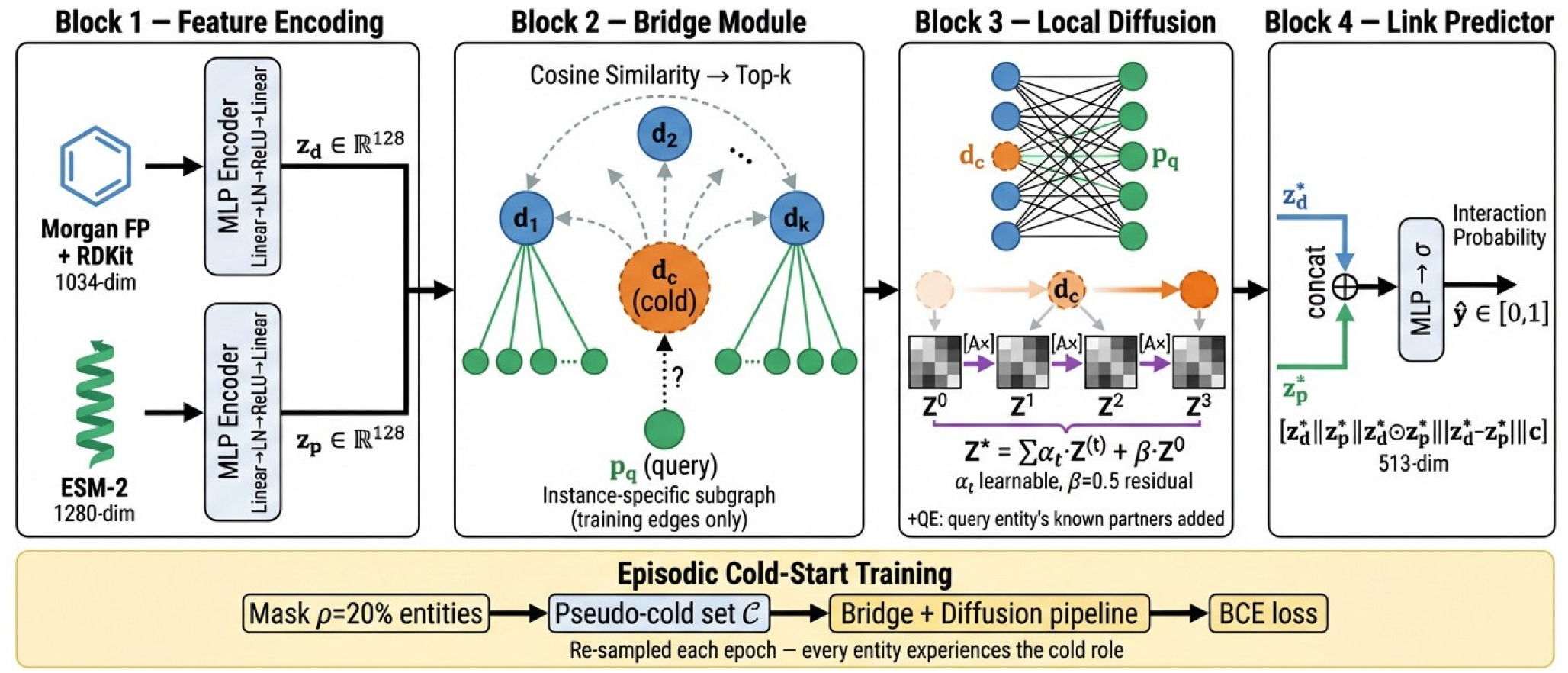
Overview of EBD-DTI. **Block 1:** Drug and protein features are independently encoded into a shared 128-dim embedding space. **Block 2:** For each cold entity, a learned bridge scorer selects the top-*k* most similar seen entities and assembles an instance-specific local subgraph including their known interaction partners. **Block 3:** *T* =3 hops of local diffusion propagate relational context from bridge neighbors to the cold entity. **Block 4:** A predictor scores drug-protein pairs from diffused embeddings. Episodic training (bottom) randomly masks 20% of entities per epoch to provide supervised gradient signal to the bridge under simulated cold-start conditions.

*Node Features*. Each drug is represented by a 1034-dimensional vector: a 1024-bit Morgan fingerprint [16] (radius=2) concatenated with 10 RDKit [11] physicochemical descriptors (molecular weight, LogP, TPSA, H-bond counts, ring statistics). Each protein is encoded as a 1280-dimensional embedding using ESM2 [13] (650M parameters) with mean pooling. Both modalities are projected into a shared *d*_*e*_-dimensional space via two-layer MLPs with LayerNorm, ReLU, and dropout:

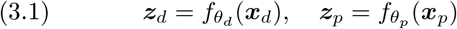

The encoder is inductive, enabling zero-shot inference for unseen entities.

#### Bridge Construction

A cold entity has no edges in the training graph. The bridge module retrieves the top- *k* nearest seen entities by cosine similarity in embedding space:

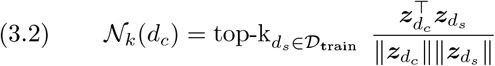

Softmax-normalized similarities define bridge weights *w*_*s*_ *∝* exp(sim(*d*_*c*_, *d*_*s*_)). The local subgraph*G*_local_ is assembled by adding the cold entity, its *k* bridge neighbors with weighted edges, and all known interaction partners of these neighbors from*Y*_train_. The resulting subgraph is instance-specific and uses only training edges, ensuring strict zero-shot inference.

#### Local Diffusion

The adjacency matrix ***A***_local_ is constructed as follows. The subgraph is bipartite (no self-loops): edges connect only drugs to proteins. Bridge edges from *d*_*c*_ to each neighbor *d*_*s*_ carry weight *w*_*s*_; all other edges (known DTI pairs) carry unit weight. Rows are *l*_1_-normalized (row-sum = 1) to obtain ***A***_local_, giving a degree-aware, asymmetric operator with no nonlinearity. No self-loops are added, so the cold entity’s initial features are preserved only through the explicit residual term *β****Z***^(0)^. Information is propagated via *T* diffusion steps:

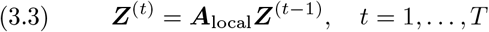

The final representation fuses all hop outputs with a residual connection:

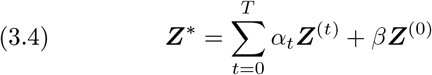

where *α*_*t*_ are softmax-normalized learnable hop weights and *β* is a fixed residual weight. Given the bipartite DTI structure (diameter 2–3 hops), *T* =3 suffices to cover the full subgraph.

### 3.4 Episodic Cold-Start Training

The bridge module receives no gradient signal under standard training, as all entities are observed. We address this with **episodic cold-start training**, which simulates cold-start conditions within each epoch.

At each epoch, a fraction *ρ* of entities is uniformly sampled as a pseudo-cold set:

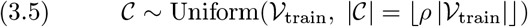

Their interactions are removed from the edge cache, and they are treated as unseen. Pairs involving pseudo-cold entities pass through the full bridge pipeline, providing direct supervision to the bridge. Support-only pairs follow a standard forward pass. Resampling each epoch ensures every entity experiences the cold role across training.

Unlike self-supervised masking (e.g., GraphMAE [6]), our approach uses supervised DTI labels, directly aligning training with the inference objective.

### 3.5 Optional Extensions

#### Query Edges (QE)

In the cold-drug setting, the query protein *p*_*q*_ appears in *G* _local_ without incident edges (only the candidate interaction to predict), leaving its post-diffusion representation close to its initial encoding. To enrich this representation, we optionally add its known training interactions to the subgraph:

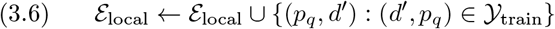

An analogous extension applies in cold-target mode for the query drug *d*_*q*_. Since the query entity is always a seen entity, QE draws only from *Y*_train_ and introduces no test-set information. We report results both with and without QE.

### 3.6 Training and Prediction

#### Training Procedure

The model is trained end-to-end with binary cross-entropy loss:

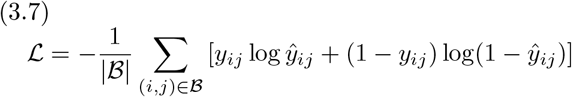

using Adam [8] (lr=10^*−*3^) with early stopping based on validation AUC (patience 10, max 60 epochs).

#### Link Prediction

For a pair (*d*_*i*_, *p*_*j*_), we compute the interaction score from post-diffusion embeddings:

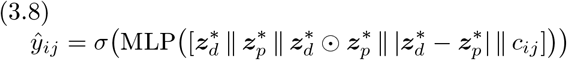

where *c*_*ij*_ is the maximum bridge weight for the cold entity. The MLP maps from 4*d*_*e*_ + 1 to 1 through a hidden layer with ReLU and dropout.

#### Cold-start evaluation protocol

We hold out 20% of entities as the cold test set ( *D*_train_ *∩ D*_test_ = ∅ for cold-drug; *P*_train_*∩ P*_test_ = ∅ for cold-target), with all their interactions removed from the training graph. Validation pairs are sampled from training-entity interactions for early stopping. We report mean *±* std over 3 seeds (520, 521, 522).

## 4. Experiments

### 4.1 Datasets

We evaluate on three widely-used DTI benchmarks (Table 1).

**Table 1.**
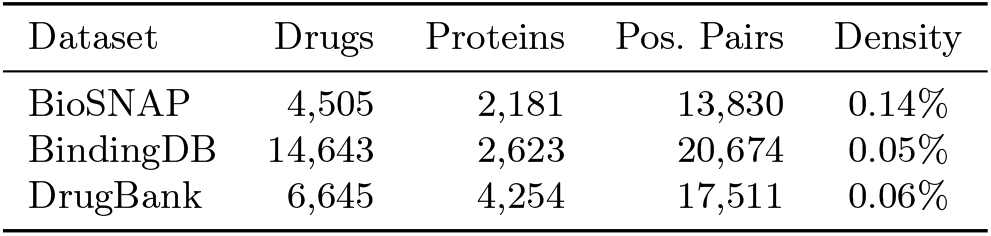
Dataset statistics.

**BioSNAP** [25] contains experimentally validated interactions from multiple sources. **BindingDB** [14] is the largest dataset with high-throughput screening interactions. **DrugBank** [20] provides curated drug– target pairs with diverse proteins.

### 4.2 Baselines

We compare against two categories of baselines.

#### Sequence-based methods

MolTrans [7], HyperAttentionDTI [24], DrugBAN [1], MlanDTI [21], and Cold-startCPI [23]. These methods are re-run on our cold-start splits using their original input features and architectures as described in their respective papers.

#### Graph-based methods

GCN [9], GraphSAGE [5], GAT [17], and APPNP [10]. All four GNN baselines are re-run on our cold-start entity splits using the *same* input features as EBD-DTI (1034-dim Morgan FP + ESM-2 embeddings), with all test-entity edges removed and no KNN augmentation, ensuring a fair zero-shot comparison under identical feature conditions.

Tables 2 and 3 report cold-drug and cold-target results (**Ours** = full EBD-DTI with Query Edges).

**Table 2.**
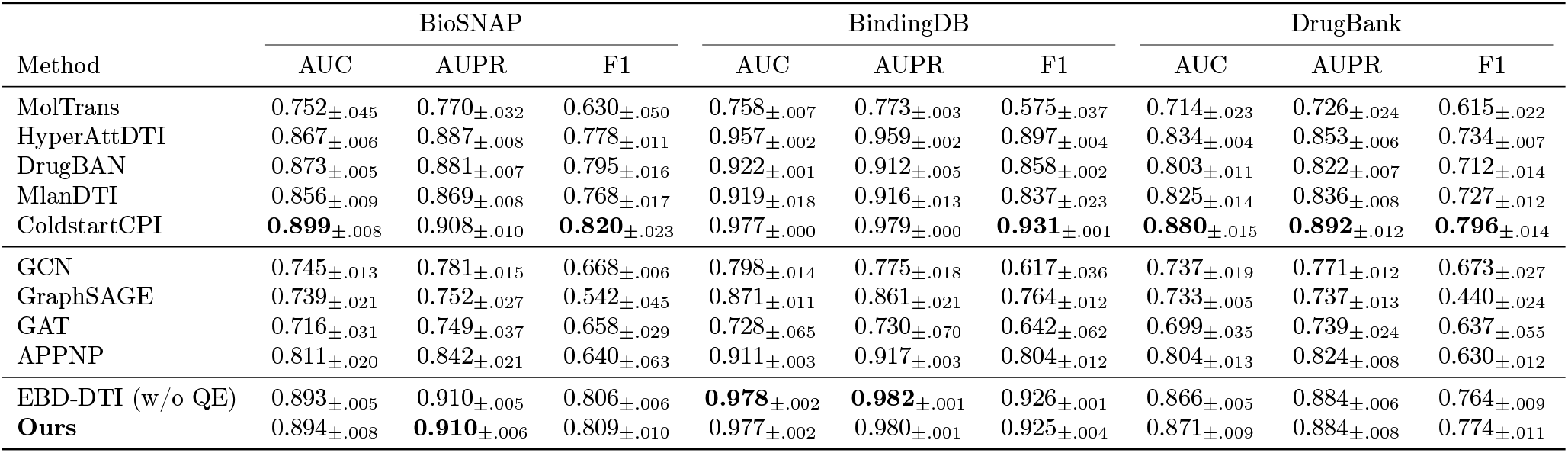
Cold-Drug results (mean*±*std over 3 seeds). **Bold**: best per column.

| Method | BioSNAP |  |  | BindingDB |  |  | DrugBank |  |  |
| --- | --- | --- | --- | --- | --- | --- | --- | --- | --- |
|  | AUC | AUPR | F1 | AUC | AUPR | F1 | AUC | AUPR | F1 |
| MolTrans | 0.752±.045 | 0.770±.032 | 0.630±.050 | 0.758±.007 | 0.773±.003 | 0.575±.037 | 0.714±.023 | 0.726±.024 | 0.615±.022 |
| HyperAttDTI | 0.867±.006 | 0.887±.008 | 0.778±.011 | 0.957±.002 | 0.959±.002 | 0.897±.004 | 0.834±.004 | 0.853±.006 | 0.734±.007 |
| DrugBAN | 0.873±.005 | 0.881±.007 | 0.795±.016 | 0.922±.001 | 0.912±.005 | 0.858±.002 | 0.803±.011 | 0.822±.007 | 0.712±.014 |
| MlanDTI | 0.856±.009 | 0.869±.008 | 0.768±.017 | 0.919±.018 | 0.916±.013 | 0.837±.023 | 0.825±.014 | 0.836±.008 | 0.727±.012 |
| ColdstartCPI | **0.899**±.008 | 0.908±.010 | **0.820**±.023 | 0.977±.000 | 0.979±.000 | **0.931**±.001 | **0.880**±.015 | **0.892**±.012 | **0.796**±.014 |
| GCN | 0.745±.013 | 0.781±.015 | 0.668±.006 | 0.798±.014 | 0.775±.018 | 0.617±.036 | 0.737±.019 | 0.771±.012 | 0.673±.027 |
| GraphSAGE | 0.739±.021 | 0.752±.027 | 0.542±.045 | 0.871±.011 | 0.861±.021 | 0.764±.012 | 0.733±.005 | 0.737±.013 | 0.440±.024 |
| GAT | 0.716±.031 | 0.749±.037 | 0.658±.029 | 0.728±.065 | 0.730±.070 | 0.642±.062 | 0.699±.035 | 0.739±.024 | 0.637±.055 |
| APNP | 0.811±.020 | 0.842±.021 | 0.640±.063 | 0.911±.003 | 0.917±.003 | 0.804±.012 | 0.804±.013 | 0.824±.008 | 0.630±.012 |
| EBD-DTI (w/o QE) | 0.893±.005 | 0.910±.005 | 0.806±.006 | **0.978**±.002 | **0.982**±.001 | 0.926±.001 | 0.866±.005 | 0.884±.006 | 0.764±.009 |
| **Ours** | 0.894±.008 | **0.910**±.006 | 0.809±.010 | 0.977±.002 | 0.980±.001 | 0.925±.004 | 0.871±.009 | 0.884±.008 | 0.774±.011 |

**Table 3.** Cold-Target results (mean*±*std over 3 seeds). **Bold**: best per column.

| Method | BioSNAP |  |  | BindingDB |  |  | DrugBank |  |  |
| --- | --- | --- | --- | --- | --- | --- | --- | --- | --- |
|  | AUC | AUPR | F1 | AUC | AUPR | F1 | AUC | AUPR | F1 |
| MolTrans | 0.694±.032 | 0.734±.039 | 0.608±.034 | 0.620±.044 | 0.627±.018 | 0.420±.001 | 0.712±.021 | 0.719±.037 | 0.585±.039 |
| HyperAttDTI | 0.783±.022 | 0.826±.027 | 0.679±.033 | 0.689±.051 | 0.705±.028 | 0.445±.055 | 0.763±.007 | 0.802±.011 | 0.648±.017 |
| DrugBAN | 0.764±.033 | 0.712±.056 | 0.652±.035 | 0.600±.043 | 0.595±.017 | 0.342±.031 | 0.719±.010 | 0.757±.005 | 0.596±.016 |
| MlanDTI | 0.848±.002 | 0.880±.006 | 0.770±.019 | 0.746±.039 | 0.754±.027 | 0.534±.061 | 0.812±.008 | 0.833±.004 | 0.699±.015 |
| ColdstartCPI | 0.876±.030 | 0.894±.036 | 0.798±.044 | **0.881**±.024 | **0.894**±.011 | **0.754**±.023 | **0.873**±.010 | **0.888**±.013 | **0.779**±.014 |
| GCN | 0.713±.033 | 0.764±.043 | 0.664±.027 | 0.425±.035 | 0.521±.019 | 0.129±.039 | 0.652±.018 | 0.701±.010 | 0.607±.019 |
| GraphSAGE | 0.737±.023 | 0.770±.028 | 0.511±.098 | 0.580±.025 | 0.581±.022 | 0.198±.027 | 0.662±.017 | 0.708±.011 | 0.500±.043 |
| GAT | 0.718±.033 | 0.753±.026 | 0.680±.038 | 0.517±.049 | 0.566±.028 | 0.327±.139 | 0.670±.024 | 0.708±.008 | 0.652±.024 |
| APNP | 0.787±.021 | 0.832±.028 | 0.607±.037 | 0.537±.048 | 0.602±.028 | 0.067±.026 | 0.713±.023 | 0.762±.015 | 0.503±.027 |
| EBD-DTI (w/o QE) | 0.882±.005 | 0.908±.008 | 0.798±.013 | 0.816±.014 | 0.827±.013 | 0.566±.015 | 0.816±.005 | 0.839±.008 | 0.702±.012 |
| **Ours** | **0.901**±.003 | **0.919**±.009 | **0.816**±.015 | 0.850±.014 | 0.850±.013 | 0.562±.020 | 0.830±.009 | 0.844±.011 | 0.707±.006 |

### 4.3 Implementation Details

EBD-DTI uses embedding dimension *d*_*e*_ = 128, bridge top-*k*=10, diffusion steps *T* = 3, residual weight *β*=0.5, batch size 256, learning rate 10^*−*3^ (Adam [8]), dropout 0.1, and pseudo-cold ratio *ρ* = 0.2. Training uses early stopping (patience 10, max 60 epochs) based on validation AUC. Drug features are 1034-dimensional (Morgan fingerprint + RDKit descriptors), and protein features are 1280-dimensional ESM-2 embeddings. Cold-start splits hold out 20% of entities as the test set (strictly zero-shot: no training edges); the remaining 80% form the training set, with a validation subset sampled from training-entity pairs for early stopping. We report mean*±* std over 3 seeds. AUC and AUPR are the primary metrics. Code is available at https://github.com/LIUYellowBlack/EBD-DTI.

#### Negative sampling strategy

Following standard practice in DTI benchmarking, we use a 1:1 positive-to-negative ratio for training, validation, and testing. Negative pairs are sampled from entity-disjoint cold-start splits (no cold entity appears in training). All methods are evaluated on the same balanced test set for fair comparison.

### 4.4 Cold-Start Results

On cold-drug (Table 2), EBD-DTI surpasses all GNN baselines, outperforming the strongest GNN (APPNP) by +10.2% AUC on BioSNAP and +7.2% on BindingDB, and achieves performance competitive with the best sequence-based method (ColdstartCPI). On cold-target (Table 3), EBD-DTI achieves the best result on BioSNAP (AUPR 0.919), while Query Edges improve AUC by +0.034 on BindingDB and +0.014 on DrugBank over the no-QE baseline. These results indicate that episodic training improves generalization under strict cold-start settings.

### 4.5 Ablation Study

Figure 2 and Table 4 present ablation results on a representative split.

**Table 4.** Ablation study (AUC, mean*±*std over 3 seeds, single split). QE: Query Edges enrichment.

| Config | BioSNAP |  | BindingDB |  | DrugBank |  |
| --- | --- | --- | --- | --- | --- | --- |
|  | CD | CT | CD | CT | CD | CT |
| **Ours (Full)** | **0.894**±.008 | **0.901**±.003 | 0.977±.002 | **0.850**±.014 | **0.871**±.009 | **0.830**±.009 |
| w/o QE | 0.893±.005 | 0.882±.005 | **0.978**±.002 | 0.816±.014 | 0.866±.005 | 0.816±.005 |
| w/o episodic | 0.876±.010 | 0.843±.019 | 0.973±.003 | 0.735±.008 | 0.852±.004 | 0.780±.022 |
| w/o bridge+diffusion | 0.689±.021 | 0.641±.018 | 0.847±.015 | 0.512±.019 | 0.698±.027 | 0.578±.024 |
| Pure features only | 0.641±.019 | 0.597±.022 | 0.782±.012 | 0.456±.021 | 0.654±.025 | 0.523±.028 |

**Figure 2.**
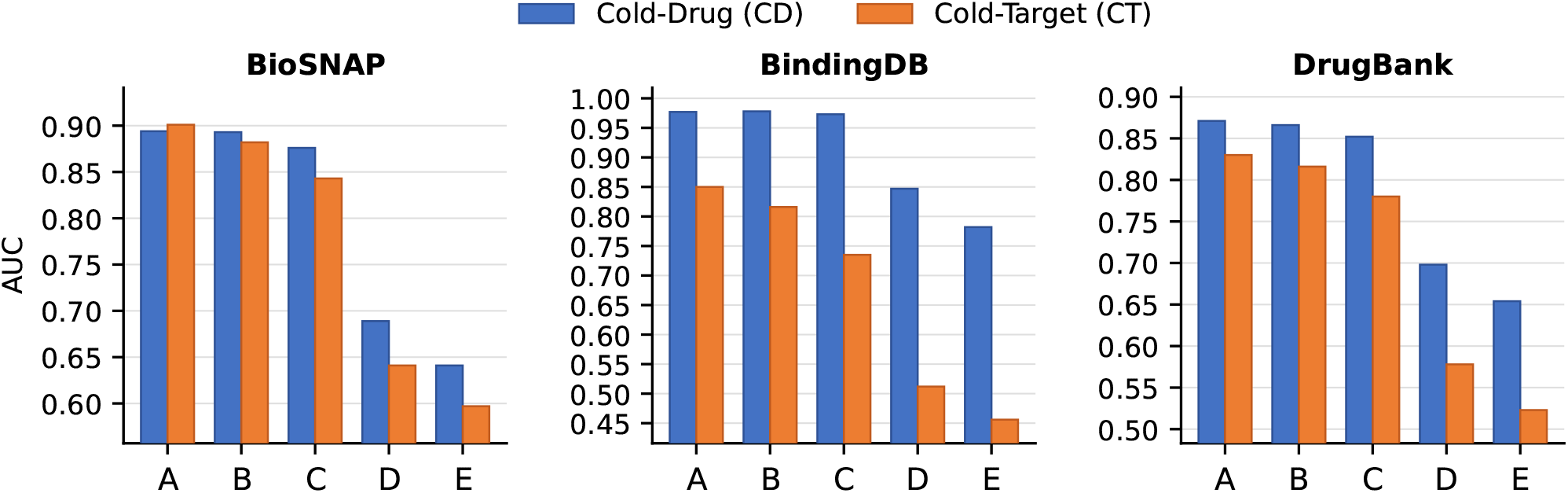
Ablation study (AUC) across three benchmarks. Blue: Cold-Drug (CD); Orange: Cold-Target (CT). **A** = Full Model, **B** = w/o Query Edges, **C** = w/o Episodic Training, **D** = w/o Bridge+Diffusion, **E** = Pure Features. The y-axis starts near each dataset’s minimum to highlight differences. Removing episodic training (C) causes the largest drop, confirming it as the dominant contribution.

Removing episodic training causes the largest drop: cold-target AUC falls from 0.901 to 0.843 on BioSNAP ( *−* 6.4%) and from 0.850 to 0.735 on BindingDB ( *−*13.5%), indicating that training signal for cold entities is the primary bottleneck. Query Edges consistently improve cold-target performance while leaving cold-drug largely unchanged, suggesting they enrich the seen entity’s context without introducing additional information. Pure feature-based models perform poorly on cold-target (AUC 0.456 on BindingDB), highlighting the importance of relational structure.

### 4.6 Discussion

*Why is cold-target harder?* The performance gap between cold-drug and cold-target reflects a feature-level asymmetry: drug molecules are small (*<*100 heavy atoms) and Morgan fingerprints capture most binding-relevant substructures, whereas protein binding depends on a localized pocket ( *∼*10–30 residues) that ESM-2’s full-sequence mean pooling dilutes. This structural mismatch makes cold-target generalization harder regardless of the graph-based enrichment applied. Query Edges partially compensate by providing relational context to the seen protein, reducing the gap by 1.4–3.4% AUC.

#### Bridge behavior

The bridge module retrieves structurally similar training entities: selected drug neighbors have substantially higher Morgan fingerprint similarity to the cold drug than random pairs, and selected protein neighbors are closer in ESM-2 embedding space than random proteins. Robustness is notable: cold-drug AUC varies within 0.005 across *k* ∈ {3, …, 30}

(Table 6), suggesting the bridge aggregates across multiple neighbors through diffusion rather than relying on a single nearest neighbor. Without episodic training, bridge neighbor selection is effectively random with respect to interaction-relevant structure; with episodic training, the encoder learns to place structurally similar and interaction-compatible entities nearby, which explains why removing episodic training causes catastrophic cold-target degradation even with the bridge architecture unchanged.

#### Query Edges: leak-free analysis

QE adds the known training-set interactions of the *query entity* (a seen entity by definition) to the local subgraph before diffusion. Since QE edges are drawn exclusively from *Y*_train_, the subgraph remains a deterministic function of training data: *G* _local_(*d*_*c*_, *p*_*q*_) = *f* (*S, Y*_train_, Encoder). No test-entity edges or neighborhoods are accessed. The asymmetric effect (Table 4: CT gains +0.019/+0.034/+0.014, CD unchanged *±* 0.005) arises because the query protein in cold-drug mode has many training partners, while the query drug in cold-target mode has fewer, limiting enrichment.

#### Computational complexity

Local diffusion ***Z***^(*t*)^ = *A*_local_***Z***^(*t−*1)^ is *O*(|ℰ _local_|*· d*_*e*_) per step, where (|ℰ |_local_ *<* 5000 and *d*_*e*_ = 128. All *T* =3 steps are negligible compared to batch gradient computation. Subgraph construction is *O*(*k* log *k* + *k ·* degree). EBD-DTI thus adds minimal overhead over a standard MLP predictor.

#### Design simplicity

We explored over a dozen alternative designs including fractional-order diffusion, curriculum masking, contrastive objectives, and hybrid prediction mechanisms, but the simplest configuration consistently performs best. This indicates that training signal alignment—not architectural complexity—is the key bottleneck for cold-start generalization.

### 4.7 Sensitivity Analysis

We report three sensitivity experiments. Table 5 sweeps the episodic mask ratio *ρ* on BioSNAP with single-drug masking. Table 6 sweeps bridge top-*k*, diffusion steps *T*, and residual weight *β* independently on BioSNAP (3 seeds, QE enabled), one parameter at a time with others fixed at defaults (*k*=10, *T* =3, *β*=0.5). Table 7 compares four episodic training strategies on a unified split (20% drugs and 20% proteins held out simultaneously).

**Table 5.** Mask ratio *ρ* sensitivity (BioSNAP, single-drug, mean AUC). **Bold**: best avg.

| $\rho$ | CD | CT | Avg |
| --- | --- | --- | --- |
| 0.05 | 0.886 | 0.869 | 0.877 |
| 0.10 | 0.887 | 0.878 | 0.883 |
| 0.15 | 0.884 | 0.864 | 0.874 |
| **0.20** | **0.893** | **0.880** | **0.886** |
| 0.25 | 0.893 | 0.876 | 0.884 |
| 0.30 | 0.895 | 0.875 | 0.885 |
| 0.35 | 0.889 | 0.875 | 0.882 |
| 0.40 | 0.895 | 0.872 | 0.883 |

**Table 6.** Bridge *k*, diffusion *T*, residual *β* sensitivity on BioSNAP (3 seeds). Format: CD*±* std / CT *±* std. **Bold**: best. Defaults *k*=10, *T* =3, *β*=0.5.

| Param | Val | CD±std / CT±std |
| --- | --- | --- |
| top-$k$ | 3 | **0.894**±.007 / 0.894±.008 |
| | 5 | 0.896±.005 / 0.894±.003 |
| | 10 | 0.892±.007 / 0.891±.008 |
| | 15 | 0.893±.008 / 0.892±.011 |
| | 20 | 0.891±.010 / 0.897±.011 |
| | 30 | 0.893±.006 / **0.899**±.007 |
| $T$ | 1 | 0.889±.006 / 0.884±.008 |
| | 2 | 0.892±.004 / 0.889±.008 |
| | 3 | 0.890±.002 / 0.894±.004 |
| | 5 | **0.895**±.006 / **0.901**±.004 |
| | 7 | 0.892±.007 / 0.900±.005 |
| $\beta$ | 0.0 | 0.891±.008 / **0.909**±.003 |
| | 0.1 | 0.893±.004 / 0.898±.003 |
| | 0.3 | 0.893±.008 / 0.897±.005 |
| | 0.5 | **0.895**±.006 / 0.897±.009 |
| | 0.7 | 0.890±.008 / 0.893±.008 |
| | 1.0 | 0.894±.010 / 0.886±.006 |

**Table 7.** Episodic variants (AUC±std, 3 seeds, unified split). **Bold**: best per column per direction.

| Mode | Dir | BioSNAP | BindingDB | DrugBank |
| --- | --- | --- | --- | --- |
| s-drug | CD | 0.885±.014 | **0.974**±.003 | **0.866**±.020 |
| | CT | 0.871±.010 | 0.819±.018 | 0.813±.010 |
| s-target | CD | 0.882±.015 | 0.969±.004 | 0.852±.019 |
| | CT | **0.884**±.012 | **0.835**±.006 | **0.822**±.009 |
| both | CD | **0.887**±.014 | 0.972±.003 | 0.861±.015 |
| | CT | 0.876±.003 | 0.832±.023 | 0.814±.021 |
| alternate | CD | 0.887±.009 | 0.972±.002 | 0.856±.019 |
| | CT | 0.878±.005 | 0.808±.027 | 0.815±.009 |

*Mask ratio (Table 5). ρ*=0.20 achieves the best average AUC (0.886) with total variation below 1.2% across *ρ* ∈ {0.05, …, 0.40 }, indicating that any moderate masking level provides sufficient cold-start signal without over-constraining the support set.

*Episodic variants (Table 7)*. Table 7 compares four masking strategies: *s-drug* (single_drug, masks only drugs), *s-target* (single_target, masks only proteins), *both* (masks both simultaneously), and *alternate* (alternates between the two). Single-mode masking dominates its respective direction: *s-target* leads on cold-target (BioSNAP CT: 0.884, BindingDB CT: 0.835), while *both* best balances CD+CT jointly, suggesting that training distribution alignment matters more than masking complexity; for unified deployment the *both* mode is preferred.

*Bridge k, diffusion T, residual β (Table 6)*. Cold-drug AUC varies by *<*0.005 across *k* ∈ {3, …, 30}, consistent with the bridge aggregating relational context from multiple neighbors through diffusion. *T* =3 matches the local subgraph diameter; *T* =5 adds only +0.007 CT at marginal additional cost. *β*=0.5 is optimal for cold-drug while *β*=0.0 is optimal for cold-target, consistent with the drug/protein feature asymmetry discussed above. Overall, all three parameters show variation below 0.025 across practical ranges, suggesting EBD-DTI does not require careful tuning.

## 5 Conclusion

We show that cold-start drug– target interaction prediction can be effectively addressed by explicitly training the bridge mechanism under the conditions encountered at test time. Standard DTI training assumes all entities are observed, creating a training–test mismatch where models are not optimized for unseen entities. EBD-DTI resolves this through episodic cold-start training, which simulates cold scenarios by masking entities during training and provides direct supervision for bridge-based inference.

This eliminates the need for test-time support sets or pre-defined graph structures.

Beyond DTI, this principle may extend to other cold-start settings where models must connect unseen entities to observed data without explicit training signal, such as recommendation systems and sparse link prediction tasks.

Empirically, episodic training is the primary driver of performance under strict zero-shot evaluation. Ablation results show that removing episodic training leads to substantial degradation (e.g., BindingDB cold-target AUC drops from 0.850 to 0.735, *−*13.5%), whereas variations in architectural components have comparatively smaller effects. This suggests that appropriate training signal, rather than model complexity, is critical for cold-start generalization. Under strict zero-shot protocols, EBD-DTI achieves strong performance across benchmarks, including leading cold-target AUPR on BioSNAP (0.919) and competitive results on BindingDB and DrugBank.

Future work includes incorporating structure-aware protein representations (e.g., binding pockets), extending episodic training to other molecular prediction tasks, and exploring adaptive masking strategies.

Overall, our results highlight the importance of aligning training with test-time conditions for reliable cold-start prediction.

## 6 Limitations

### Human dataset

The Human DTI benchmark (2,001 proteins, 6,728 pairs) was excluded from the main evaluation due to high variance under strict cold-start splits. With only 2,001 proteins, a 20% cold-target holdout leaves fewer than 400 test proteins; combined with sparse interactions, performance estimates across seeds vary by up to *±* 0.05 AUC—making comparisons unreliable. Under this unstable regime, GCN achieves 0.788 AUC while EBD-DTI achieves 0.766 (cold-target), a difference within the noise floor. We include Human results in preliminary experiments but omit it from the main table to avoid misleading conclusions.

### ESM-2 protein features

EBD-DTI encodes proteins using mean-pooled ESM-2 embeddings over the full sequence. Binding, however, depends on a localized pocket ( *∼* 10–30 residues), and full-sequence pooling dilutes pocket-relevant signal. This structural mismatch likely explains the systematic cold-target gap relative to cold-drug performance, as Morgan fingerprints for drugs more faithfully capture the substructures responsible for binding. Integrating pocket-aware protein representations (e.g., from AlphaFold2 structures) is a natural extension.

### Inductive generalization scope

EBD-DTI is designed for the *single-entity* cold-start setting (cold-drug or cold-target separately). The fully inductive *blind* setting—where both the drug and the protein are unseen simultaneously—is substantially harder and not addressed here. Extending episodic training to blind cold-start (masking drug–protein pairs jointly) is left for future work.

## 7 Exploration of Alternative Designs

We systematically evaluated a wide range of alternative designs before arriving at the final EBD-DTI formulation. These include: (i) family-aware bridge selection based on protein clustering; (ii) curriculum-based masking strategies (degree-based and confidence-based); (iii) auxiliary reconstruction and contrastive objectives (e.g., SBO and InfoNCE); (iv) hybrid prediction mechanisms combining bridge and content features; (v) post-hoc label propagation; and (vi) more complex diffusion variants such as fractional-order and gradient-based propagation.

Across all experiments, these alternatives consistently failed to improve performance over the proposed design, and often degraded cold-target results. Auxiliary objectives and curriculum strategies introduced instability or reduced training diversity, while more complex diffusion mechanisms led to over-smoothing on small local subgraphs. Hybrid and post-hoc strategies were ineffective due to weak or uninformative confidence signals.

These results suggest that the primary challenge in cold-start DTI is not architectural complexity, but the availability of appropriate training signal. The simplicity of EBD-DTI—combining episodic training with local subgraph diffusion—appears sufficient to capture the essential structure required for generalization.

## Acknowledgments

This work was supported by the National Natural Science Foundation of China and the Fundamental Research Funds for the Central Universities, Nankai University.

